# Human colonization with *Phytobacter* co-harboring *bla*_IMP-4_ and *mcr-9.1* highlights its potential as emerging human pathogen

**DOI:** 10.1101/2025.10.09.681365

**Authors:** Anna Tumeo, Aneta Kovarova, Francesca McDonagh, Kate Ryan, Christina Clarke, Georgios Miliotis

## Abstract

*Phytobacter* is a recently delineated, frequently misidentified genus within the order Enterobacterales. Following two rare cases of patient colonization with multidrug resistant *Phytobacter* in Ireland, this study presents a genus-wide genomic analysis that aims to define the pathogenic potential of *Phytobacter* species, with emphasis on their role as emerging human pathogens and reservoirs of carbapenemases. Two carbapenemase-encoding isolates were recovered from rectal swabs in Ireland in 2024 and were initially identified as *Phytobacter* by MALDI-ToF. Whole-genome sequencing with *in silico* species typing (dDDH, ANI) provided definitive taxonomic resolution. A genus-wide maximum-likelihood core-genome phylogeny was reconstructed, and the plasmidome and resistome were bioinformatically profiled across all available *Phytobacter* genomes. Phenotypic susceptibility of the Irish isolates was determined through minimum inhibitory concentration (MIC) testing. The Irish isolates (*P. diazotrophicus* E787336 and *P. ursingii* E980862) are the first reported *Phytobacter* strains carrying both plasmid-borne *bla*_IMP-4_ and *mcr-9*.*1* in the genus. MIC testing confirmed resistance to aztreonam, aminoglycosides, cephalosporins, fluoroquinolones, and the β-lactam/β-lactamase inhibitor combination piperacillin–tazobactam. Across 34 *Phytobacter* genomes examined, 22 distinct plasmid replicon types were identified in 22 isolates, often shared across species. The genus-wide resistome encompassed 71 genes, more than half predicted to be acquired, with carbapenemases detected in 26.5% (9/34) of the genomes. In summary, *Phytobacter* harbors a diverse, plasmid-borne resistome including carbapenemases, with documented cases of human colonization and infection. These findings support its recognition as an emerging pathogen and reservoir of antimicrobial resistance, underscoring the need for improved clinical identification, genomic surveillance, and preparedness for limited treatment options.

**Author summary:** Since *Phytobacter* was first characterized in 2007, this bacterial genus has mainly been associated with plant growth promotion. More recently, however, increasing reports of human infections have raised concerns about its potential as emerging bacterial pathogen. These are further underscored by the description of multidrug resistant isolates capable to withstand different classes of antimicrobials, including carbapenems which are often used as a last line of treatment. Following the rare finding of multidrug resistant *Phytobacter* in two patients in Ireland, we combined bioinformatics and laboratory testing to characterize the antimicrobial resistance profile of this overlooked bacterial genus. Our results uncover the variety of resistance determinants, including to carbapenems, which is encoded in the genomes of *Phytobacter*. This shows its potential as hidden reservoir of drug resistance and emerging bacterial pathogen. We encourage improved clinical recognition and monitoring of *Phytobacter* to better anticipate infections with limited therapeutic options.

## 1. Main text

### 1.1 Introduction

The genus *Phytobacter* was first described in 2007 within the order Enterobacterales from the disentanglement of the *Erwinia herbicola*/*Enterobacter agglomerans* complex (1) and currently includes four validly published species: *P. ursingii, P. diazotrophicus, P. palmae*, and *P. massiliensis* (2–5). Originally strictly linked to plant growth promotion (4), *Phytobacter* has increasingly gained attention as opportunistic pathogen due to reports of human infections attributed to members of this genus (6– 10).

Due to its relatively new taxonomic classification and to the lack of specificity of routine clinical typing methods towards this genus, including MALDI-ToF mass spectrometry, *Phytobacter* has frequently misidentified and misassigned to other *Enterobacterales* (e.g., *Pantoea, Metakosakonia, Kluyvera*), particularly by routine MALDI-TOF MS (1,11,12). Although modern typing tools based on whole-genome sequencing such as digital DNA-DNA hybridization (dDDH) and average nucleotide identity (ANI) allow for accurate taxonomic resolution of *Phytobacter* strains (13,14), misidentification of *Phytobacter* remains common, especially in clinical settings, hindering our understanding of its prevalence in clinical environments. prevalence in clinical environments.

Previous genomic surveys report a broad ARG repertoire in *Phytobacter* including ESBLs, carbapenemases, and colistin resistance determinants, consistent with multidrug-resistant (MDR) phenotypes (15,16). Phenotypically, *Phytobacter* isolates have shown resistance to aminoglycosides, cephalosporins, fluoroquinolones, and fosfomycin, with acquired carbapenem resistance mediated, amongst others, by *bla*_KPC-2_ and *bla*_NDM-1_ (8,15,17,18).

Despite increasing reports implicating *Phytobacter* in human disease, its genomic features, including the plasmidome and resistome, remain poorly characterized. Here, we describe two rare cases of patient colonization with MDR *Phytobacter* in Ireland and, through a genus-wide genomic analysis, provide new insights into its emerging pathogenic and resistance potential.

### 1.2 Results

#### 1.2.1 Genome assembly and species typing

A genus-wide all-vs-all ANI comparison of *Phytobacter* genomes outlined four distinct clusters, each sharing >97% ANI (**Figure S1**), formed by the four validly published *Phytobacter* species: *P. massiliensis* (19), *P. diazotrophicus* (4), *P. palmae* (3), and *P. ursingii* (12). While both Irish isolates were originally classified as *P. ursingii* by MALDI-ToF, ANI revealed genomic divergence, suggesting their taxonomic resolution into *P. diazotrophicus* (E787336) and *P. ursingii* (E980862).

Taxonomic classification of E787336 strain as *P. diazotrophicus* and E980862 strain as *P. ursingii* was supported by respectively 98.7% and 97.5% ANI to the corresponding type strains and further confirmed by 90.7% and 81.5% dDDH values with these genomes (**Table 1**).

**Table 1.** Species typing of E787336 and E980862 strains based on ANI and dDDH compared to respectively *P. diazotrophicus* type strain DSM 17806^T^ and *P. ursingii* type strain ATCC 27989^T^. Original taxonomic classification by MALDI-ToF and corresponding confidence values are also reported.

| Comparison: | ANI | dDDH | MALDI-ToF |
| --- | --- | --- | --- |
| E787336 strain vs<br>*P. diazotrophicus* DSM 17806<sup>T</sup> | 98.7% | 90.7% [88.5 - 92.5%] | *P. ursingii* (2.09) |
| E980862 strain vs<br>*P. ursingii* ATCC 27989<sup>T</sup> | 97.5% | 81.5% [78.6 - 84.1%] | *P. ursingii* (2.27) |

#### 1.2.2 Phylogenomic analysis

Bootstrap-supported core-genome phylogeny confirmed clustering of E787336 strain within the same clade as all other *P. diazotrophicus* genomes, although this strain showed divergence from the main species lineage (**Figure 1**). Particularly, E787336 strain formed a monophyletic clade with *Phytobacter sp. MRY16-398*, originally classified under the genus *Metakosakonia*, isolated from ascites of a patient in Japan in 2015. Except for *P. ursingii* PSU_26, E980862 strain clustered within a monophyletic clade formed by all *P. ursingii* genomes, the closest phylogenetically being *P. ursingii* reference genome CAV1151, a clinical strain isolated from a human host in Virginia, USA, in 2009. SNP analysis revealed 578 core SNPs between E787336 strain and *Phytobacter sp. MRY16-398* (**Table S2**), while 2201 core SNPs were identified between E980862 and CAV1151 strains (**Table S2-S3**). Given an estimated evolution rate of *Phytobacter* equal to 1.85E-6 substitutions/site/year (16), these distances are incompatible with recent transmission, therefore support unrelated acquisitions.

**Figure 1:**
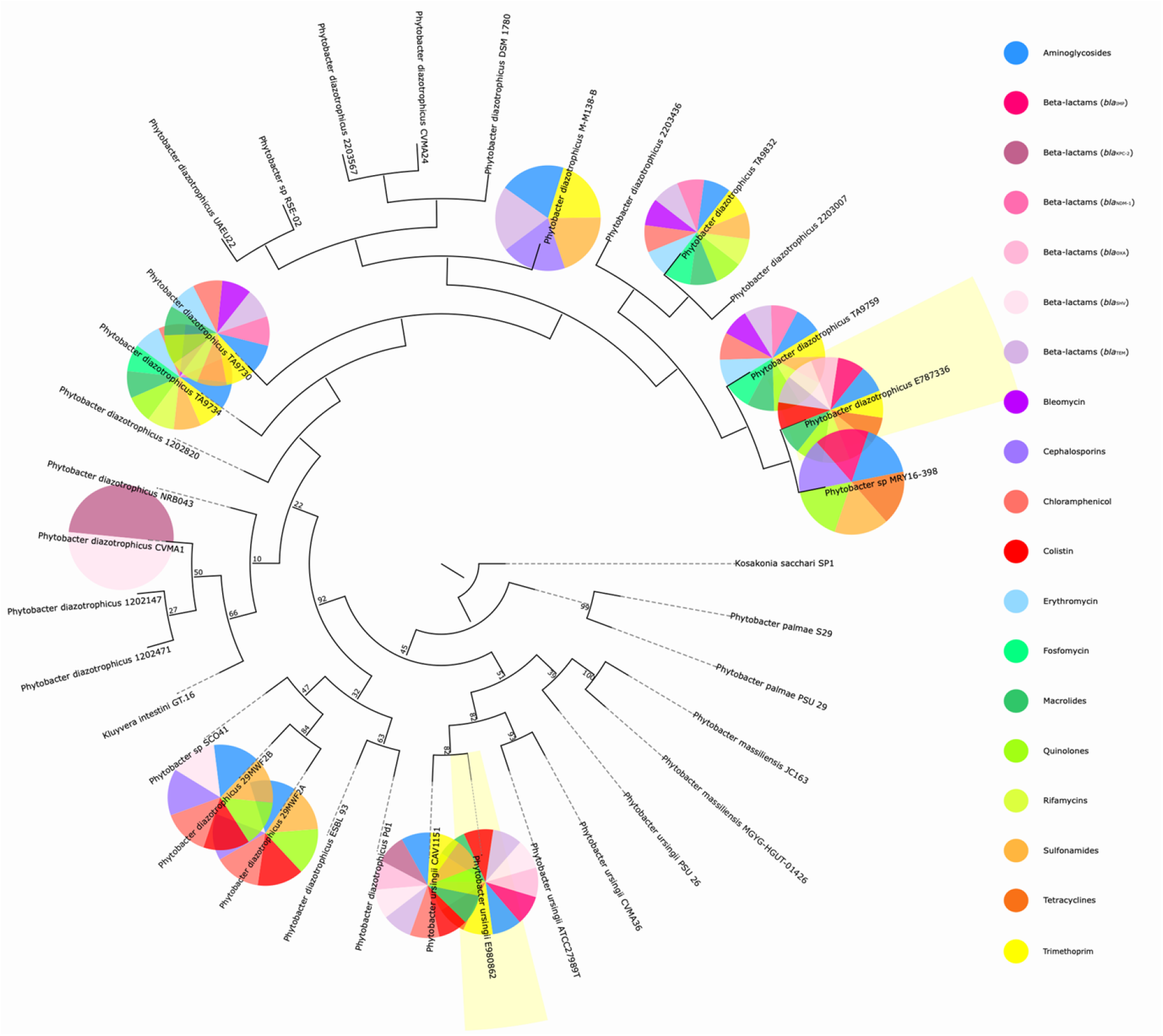
Bootstrap-supported core-genome phylogeny of *Phytobacter* genus. *Kosakonia sacchari* SP1 strain is included as an outgroup. E787336 and E980862 strains are highlighted in yellow. For each genome, predicted acquired ARGs are depicted in the accompanying pie charts.

#### 1.2.3 Genus-wide characterization of the plasmidome and resistome of *Phytobacter*

Prediction of plasmid sequences from draft genome assemblies led to the identification of two conjugative plasmids shared by the Irish isolates: a multi-replicon (IncFIB/IncFIC) plasmid (AA831) predicted to originate from *Phytobacter sp. MRY16-398*; and an IncHI2A plasmid (AA739) with predicted origin in *P. ursingii*. Two additional mobilizable plasmid replicons (AA519 and AB040) were predicted in E787336 strain. Two mobilizable (AB043 and AB050) and two non-mobilizable (AA113 and AB536) plasmid replicons were also predicted in E980862 strain.

At the whole-genus level, 22 distinct and inter-species plasmid types were identified across nearly 65% (22/34) of the investigated genomes (**Table S4**). The multi-replicon IncFIB/IncFIC plasmid was the most ubiquitously detected plasmid type, shared by >80% (18/22) of these genomes.

A single conjugative IncHI2A plasmid (~300 kb) (AA739) carried all predicted acquired ARGs in both E787336 (n=16) and E980862 (n=14) strains. These included resistance genes to aminoglycosides, beta-lactams, cephalosporins, macrolides, sulfonamide, tetracycline, trimethoprim, and to both classes of last-resort antibiotics carbapenems (*bla*_IMP-4_) and colistin (*mcr-9*.*1*). In E787336 strain, AA739 also carried *qnrB20*, potentially conferring resistance to fluoroquinolones. The Irish isolates represent the only cases of co-occurring plasmid-borne *bla*_IMP-4_ and *mcr-9*.*1* in the genus.

A comprehensive search for ARGs in all investigated genomes of *Phytobacter* led to the identification of a genus-wide resistome comprising 71 ARGs (**Figure 2**). Among these, eight chromosomal ARGs including *acrB, marA, ompA, emrR, msbA, CRP, H-NS*, and *cpxA* appeared to be intrinsic to the genus *Phytobacter* as these were detected in all investigated genomes (34/34) (**Figure 2A**). Additionally, both components of the OqxAB efflux pump, conferring resistance to multiple agents including fluoroquinolones, were chromosomally detected in the same subset of nearly 80% (27/34) of the genomes.

**Figure 2:**
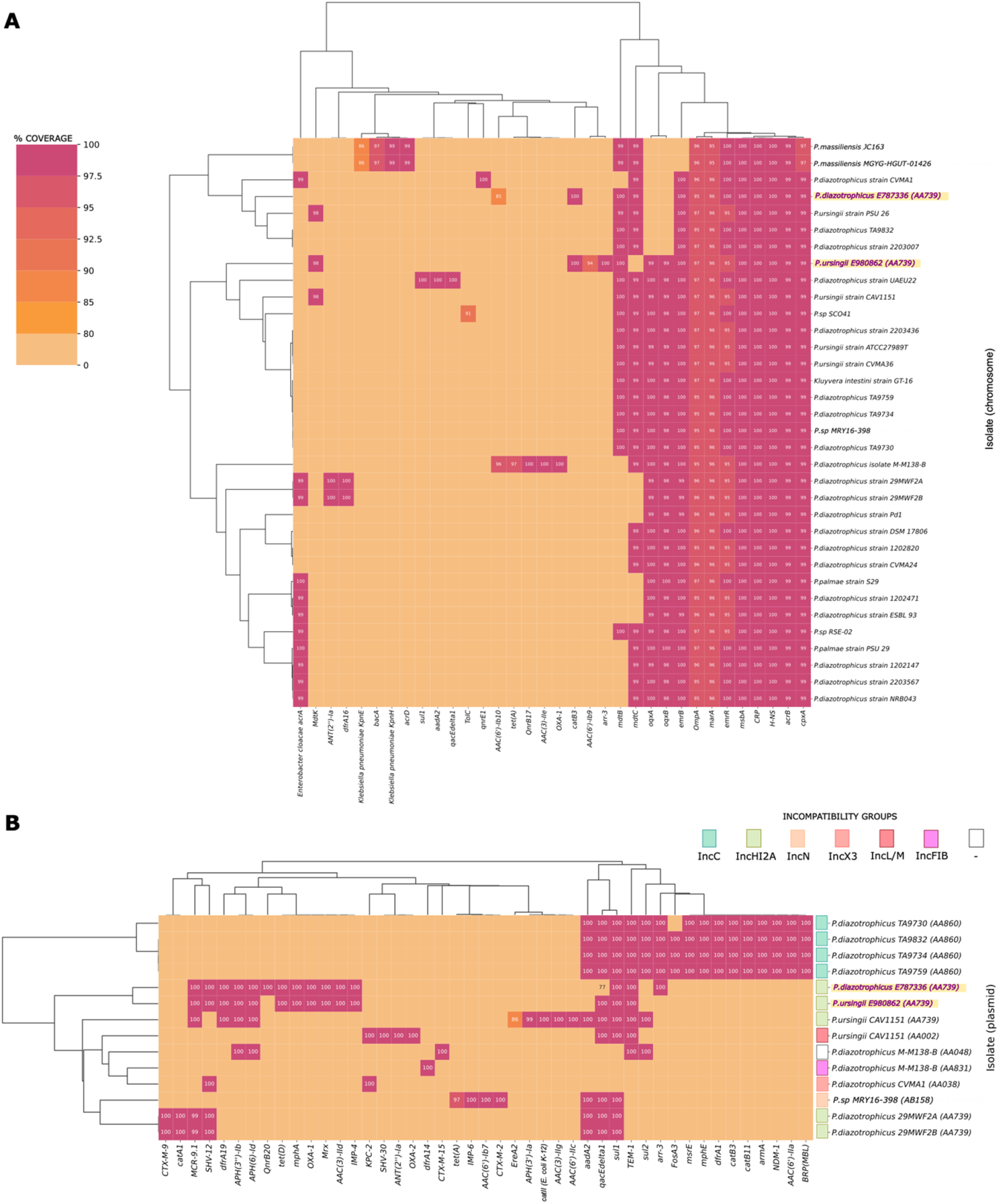
*Phytobacter*’s resistome. Distribution of antibiotic resistance genes (ARGs) identified in all *Phytobacter* genomes on both (**A**) chromosomal and (**B**) predicted plasmid sequences with color-coded associated predicted incompatibility types.

Over 50% (37/71) of the identified ARGs were exclusively predicted on plasmid sequences across 12 distinct *Phytobacter* genomes. These included carbapenemases (*bla*_IMP_, *bla*_NDM-1_, *bla*_KPC-2_) and colistin resistance determinants (*mcr-9*.*1*) which were identified in respectively 75% (9/12) and >40% (5/12) of these genomes (i.e., 26.5% and 14.7% of all *Phytobacter* genomes) (**Figure 2B**). Remaining ARGs included resistance determinants to aminoglycosides (*aac(3), aac(6), aph(3’’)*, and *aph(6)* genes), beta-lactams (*bla*_OXA_ genes and *bla*_TEM-1_), cephalosporins (*bla*_CTX-M_ and *bla*_SHV_ genes), fosfomycin (*fosA3*), fluoroquinolones (*qnrB20*), macrolides (*mph* genes and *msrE*), phenicols (*cat*-type genes), sulfonamide (*sul1* genes), rifamycin (*arr-3*), tetracycline (*tet* genes), and trimethoprim (*dfr* genes). Notably, among the most frequently detected genes, *qacEdelta1*, conferring resistance to disinfecting agents, was predicted in over 83% (10/12) of the plasmid-harboring genomes.

#### 1.2.4 MIC testing

MIC testing confirmed phenotypic resistance of E787336 and E980862 strains to a shared set of 7 out of the 32 antibiotics tested, particularly to aminoglycosides (gentamicin [Gm], tobramycin [To]), third- to fifth-generation cephalosporins (cefepime [Cpe], ceftazidime [Caz], cefotaxime [Cft], ceftolozane-tazobactam [C/T]), and the monobactam aztreonam [Azt] (**Figure 3**). Additionally, E787336 strain showed resistance to the β-lactam/β-lactamase inhibitor combination piperacillin-tazobactam (P/T) and to the fluoroquinolones ciprofloxacin (Cp) and norfloxacin (Nxn). Both E787336 and E980862 strain demonstrated sensitivity to all tested carbapenems (ertapenem, imipenem, and meropenem) and colistin. Results for colistin MIC testing using UMIC Colistin strips (Bruker Daltonics GmbH & Co. KG, 28359 Bremen, Germany) are available as **Table S5**.

**Figure 3:**
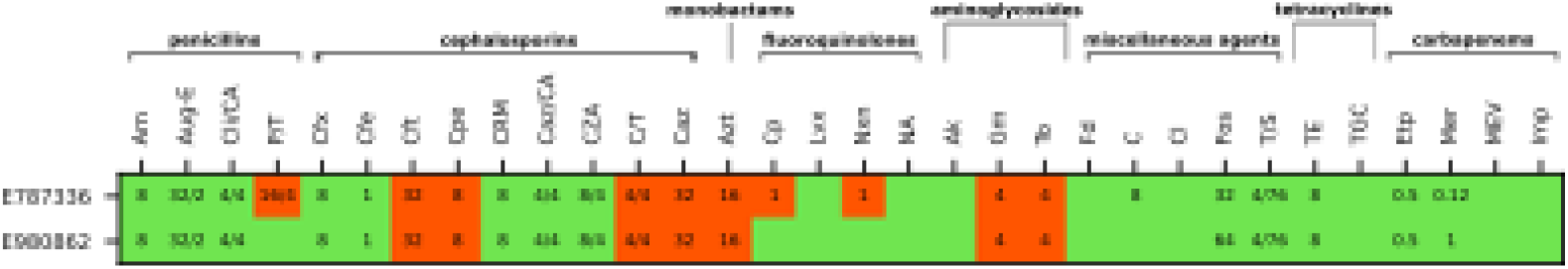
MIC testing. MIC results indicating susceptibility (green) or resistance (red) of E787336 and E980862 strains to all tested antibiotics (n=32). For each antibiotic, the observed minimum inhibitory concentration value (mg/L) is reported.

### 1.3 Discussion

Although the taxonomy of the genus *Phytobacter* has now largely been resolved, both genus-wide ANI analysis and bootstrap-supported core-genome phylogeny pinpoint residual inconsistencies. Particularly, taxa currently named under different species including *Kluyvera intestini* GT-16 strain and *Phytobacter* species MRY16-398, RSE-02, and SCO41 are likely to be reclassified to *P. diazotrophicus*. This is supported by >97% ANI of these genomes to *P. diazotrophicus* type strain DSM17806^T^ and by their clustering within *P. diazotrophicus* monophyletic clade. In addition, preliminary misclassification by MALDI-ToF of E787336 strain as *P. ursingii*, later reclassified as *P. diazotrophicus* through multifactorial *in silico* genomic species typing, underscores the persisting difficulties in the correct identification of *Phytobacter* isolates in clinical settings without the use of modern sequencing-dependent typing tools.

To date, reports of infections attributed to *Phytobacter* species worldwide remain limited (6,7,9,10). The true incidence of human infections, however, is likely underestimated due to the previously mentioned limitations in the specificity of clinical typing methods towards *Phytobacter* species. These isolates constitute the first documented cases of multidrug-resistant *Phytobacter* colonization in patients in Europe. Given these cases are colonization rather than infection; we therefore avoid inferring pathogenicity from these two observations.

A broad, mobilizable ARG profile was predicted for both Irish isolates on the same conjugative IncHI2A plasmid (AA739). IncHI2A has demonstrated clinical relevance as previously reported to carry both *mcr-9* (20) and *bla*_IMP-4_ genes (21,22). A study conducted across three Irish hospitals also highlighted its co-occurrence with *mcr-9*.*1* genes (23). Of note, AA739 was likewise found to carry *mcr-9*.*1* in E980862 phylogenetic neighbor *P. ursingii* CAV1151 strain (human source, Virginia, USA, 2009). E787336 and E980862 strains represent the first occurrence of co-carriage of acquired *mcr-9* and *bla*_IMP-4_ in the genus *Phytobacter*.

The MDR phenotype of the Irish isolates assessed through MIC testing reflected their AMR genomic profiles. Phenotypic resistance to aminoglycosides can be explained by the genomic presence of *aac(3)-IId, aph(3’’)-Ib, aph(6)-Id*, while resistance to third- to fifth-generation cephalosporins is likely mediated by *bla*_SHV-12_ and to piperacillin-tazobactam by *bla*_IMP-4_ (24). The identification of *qnrB20* in E787336 strain also explains its additional phenotypic resistance to the fluoroquinolones Cp and Nxn. While none of the identified ARGs would alone confer resistance to aztreonam, this may be induced in both Irish isolates by the combined production of the acquired broad-spectrum ß-lactamase *bla*_TEM-1_ and metallo-ß-lactamase *bla*_IMP-4_, and by the activity of chromosomal MDR efflux pumps. Collectively, these findings suggest a potential horizontal AMR acquisition pathway and constitute a valid example of the risks posed by *Phytobacter* species as MDR reservoirs and/or vectors in both clinical settings and human hosts.

Although they both harbored acquired *mcr-9*.*1* and *bla*_IMP-4_, the Irish isolates remained susceptible to colistin and carbapenems. However, while susceptibility to colistin despite the presence of *mcr-9*.*1* has previously been described (23,25,26), susceptibility to carbapenems in *bla*_IMP-4_-positive isolates is rare. A study investigating environmental contamination with carbapenemase-producing Enterobacteriaceae in Australia reported the isolation of *bla*_IMP-4_-positive isolates susceptible to imipenem and meropenem (27). Susceptibility to both carbapenems was also reported in *bla*_IMP-4_-positive clinical isolates of *Serratia marcescens, Klebsiella pneumoniae*, and *Escherichia coli* (28). Preferential susceptibility to either meropenem or imipenem is rather common in isolates harboring *bla*_IMP-1_ and *bla*_IMP-6_ genes (29,30).

At the whole-genus level, the intrinsic resistome of *Phytobacter* mainly consists of chromosomally encoded structural and/or regulatory subunits of MDR efflux pumps. While this is compatible with previous observations of MDR and XDR isolates of *Phytobacter* (15,16), not all genomes harbored all efflux systems essential components. For instance, despite the ubiquity of *acrB* and *marA*, the remaining subunit of the AcrAB efflux pump, *acrA*, was only found in 10 genomes of *Phytobacter*. Similarly, although *emrR* and *emrB* were ubiquitously found in the genus, all investigated genomes lacked the remaining subunit of the EmrAB complex, *emrA*. Additionally, the outer membrane channel TolC, necessary to most of these efflux systems to work, was detected in only one genome (*Phytobacter sp. SCO41*).

The most common plasmid in *Phytobacter* genus is the conjugative multi-replicon plasmid AA831 (IncFIB/IncFIC). Studies show that IncF plays a major role in the dissemination of AMR among *Enterobacteriaceae* (31) and can often assume a multi-replicon status (32), likely resulting in a broader and more diversified ARG content, hence in better survival under antibiotic pressure (33). In this case, no ARGs were detected on AA831 except for *dfrA14* in *P. diazotrophicus* M-M138-B strain. However, limitations in the interpretation of these results should be acknowledged, as complete plasmid sequence reconstruction was hampered by the draft status of the genome assemblies.

Despite the overall high variability in *Phytobacter*’s predicted acquired resistome, clusters of conserved ARGs across multiple species were identified for each plasmid incompatibility type, suggesting inter-species horizontal gene transfer events to have likely contributed to delineating the observed MDR profiles.

*Phytobacter* harbors a diverse acquired resistome on clinically relevant plasmids, which includes multiple carbapenemases (*bla*_IMP_, *bla*_NDM-1_, *bla*_KPC-2_) and resistance determinants against disinfecting agents (*qacEdelta1*). This further underlines the possibility of enhanced persistence of *Phytobacter* species in clinical settings, their role as carbapenemase reservoirs, and of infections with limited therapeutic options. Combined with demonstrated instances of MDR colonization in patients and documented infection cases, these results support recognition of *Phytobacter* as an emerging human pathogen and potential reservoir of carbapenemases. These findings ultimately argue for strengthened genomic characterization and clinical recognition to mitigate routine clinical misidentification of *Phytobacter* isolates, and for improved targeted surveillance to anticipate limited therapeutic options.

### 1.4 Materials and methods

#### 1.4.1. Isolates collection

Two *Phytobacter* strains were reported to Galway Reference Laboratory Service. Strains E787336 and E980862 were isolated from rectal swabs of a 77-year-old male (January 2024) and a 6-year-old female (July 2024), respectively. Both strains were originally classified as *P. ursingii* by Matrix-Assisted Laser Desorption/Ionization-Time of Flight (MALDI-ToF) mass spectrometry (MALDI Biotyper, Bruker Daltonics GmbH, Germany).

#### 1.4.2 DNA sequencing and genome assembly

DNA extraction was conducted with Qiagen EZ1 instrument and the EZ1/2 Tissue Kit (Qiagen, Tegelen, Netherlands) following manufacturer’s instructions. Library preparation was performed with the Illumina DNA Prep Kit (Illumina, Inc., San Diego, California, USA), and the Biomek 4000 instrument (Beckman Coulter Life Sciences, CA, USA). Total DNA concentration was quantified using the dsDNA High Sensitivity kit and Qubit fluorometer v1.0 (ThermoFisher Scientific, Waltham, MA). Second-generation sequencing was conducted using P1 600cycle kit and the NextSeq1000 (PE150; Illumina, Inc., San Diego, California, USA). Following quality control of raw reads (>25 Q score) using [fastp](https://github.com/OpenGene/fastp) (34), a total of respectively 5911112 (E787336) and 1427390 (E980862) filtered reads were submitted to genome assembly using Shovill v1.1.0, setting minimum contig length to 200. Assembled genomes were assessed for quality, completeness, and contamination using [quast](https://github.com/ablab/quast) v5.3.0 and [checkM](https://github.com/Ecogenomics/CheckM) v1.2.3 (35). Complete genome assembly statistics is available as **Table S1**.

#### 1.4.3 *In vitro* and *in silico* species typing

To provide a broader genomic perspective of E787336 and E980862 strains within the genus *Phytobacter*, a cluster map was generated using [ANIclustermap](https://github.com/moshi4/ANIclustermap) v1.3.0 including all canonical *Phytobacter* genomes available in NCBI GenBank database (accessed June 2025), excluding MAGs, anomalous assemblies, and suppressed genomes (n=32). Genomic species typing was then confirmed based on ANI and dDDH using the [Genome-to-Genome Distance Calculator](https://ggdc.dsmz.de/distcalc2.php) (36) against the corresponding species type strains (i.e., *P. diazotrophicus* type strain DSM 17806T and *P. ursingii* type strain ATCC 27989T).

#### 1.4.4 Phylogenomic analysis

Genome annotation was conducted with Prokka v1.14.5 (37), and [Roary](https://github.com/sanger-pathogens/Roary) v3.13.0 (38) was used to define *Phytobacter*’s pangenome and core genome. A Maximum Likelihood phylogeny of E787336 and E980862 strains was inferred based on a multi-sequence alignment of all core genes (n=523) using [RAxML](https://github.com/stamatak/standard-RAxML) v8.2.13 (39) with GTRGAMMA model and 500 bootstraps. The taxonomically adjacent *Kosakonia sacchari* SP1 was included as outgroup. Snp-dists was used to assess clonality of closely clustering genomes. The [ETE toolkit](https://etetoolkit.org/) (40) was used for three manipulation, analysis, and visualization.

#### 1.4.5 Genus-wide characterization of the plasmidome and resistome of *Phytobacter*

All genomes (n=34) were investigated for the presence of plasmidic contigs and their predicted sequences were individually reconstructed using [MOB-recon](https://github.com/phac-nml/mob-suite) (MOB-suite) (41). *Phytobacter*’s resistome was characterized running [Abricate](https://github.com/tseemann/abricate) v1.0.0 with CARD 2023 database (42) on all chromosomal and predicted plasmid sequences.

#### 1.4.5 MIC testing

Phenotypic susceptibility of E787336 and E980862 strains to 32 antibiotics was assessed through minimum inhibitory concentration (MIC) testing with MicroScan Gram-negative 63 Panel (Beckman Coulter, CA, USA) following the manufacturer’s instructions. Primary interpretation used EUCAST v15.0 (2025) breakpoints. In accordance with both EUCAST and CLSI guidelines, *Escherichia coli* ATCC 25922 and *Pseudomonas aeruginosa* ATCC 27853 were included as controls. Susceptibility to increasing concentrations (range: 0.0625 – 64 mg/L) of colistin was confirmed through MIC testing using UMIC Colistin strips (Bruker Daltonics GmbH & Co. KG, 28359 Bremen, Germany) following the manufacturer’s instructions. The colistin-susceptible *E. coli* ATCC 25922 and resistant *E. coli* NCTC 13846 (*mcr-1* positive) were included as reference strains.

## 2. Acknowledgments

The authors acknowledge the Microbiology Department at University Hospital Galway for facilitating access to MALDI-TOF MS.

## 3. Supporting information

**Figure S1:** (Figure S1) **Genus-wide average nucleotide identity analysis**. Color map showing clustering of *Phytobacter* genomes based on average nucleotide identity. E787336 and E980862 strains are highlighted in yellow.

**Table S1:** (Document S1) **Genome assembly statistics and accession numbers for E787336 and E980862 strains**.

**Table S2:** (Document S1) **Core SNP analysis conducted on *P. diazotrophicus* genomes**.

**Table S3:** (Document S1) **Core SNP analysis conducted on *P. ursingii* genomes**.

**Table S4:** (Document S2) **Prediction of plasmid sequences from *Phytobacter* genome assemblies**. Details are reported for each plasmid sequence including correspondent number of contigs, size, GC content, predicted replicon and relaxase type(s) and accession(s), predicted mobility, mash neighboring, clustering, and host range.

**Table S5:** (Document S1) **MIC testing using UMIC Colistin strips (Bruker Daltonics GmbH & Co. KG, 28359 Bremen, Germany)**. Increasing concentrations of colistin were tested (0.0625-64 mg/L). The colistin-susceptible *E. coli* ATCC 25922 and resistant *E. coli* NCTC 13846 (*mcr-1* positive) were included as reference strains. **\***GC: growth control.

## 5. Data reporting

The draft genome sequence of *P. diazotrophicus* E787336 and *P. ursingii* E980862 are available in GenBank database under accession numbers SAMN47937952 and SAMN47937953, BioProject PRJNA1250442.

## 7. Author contributions

**Anna Tumeo**: Conceptualization, Methodology, Software, Formal analysis, Data Curation, Visualization, Writing-Original Draft, Writing – Review & Editing. **Aneta Kovarova**: Investigation, Writing – Review & Editing. **Francesca McDonagh**: Investigation, Writing – Review & Editing. **Kate Ryan**: Investigation, Writing – Review & Editing. **Christina Clarke**: Resources, Writing – Review & Editing. **Georgios Miliotis**: Conceptualization, Methodology, Writing – Review & Editing, Supervision, Funding acquisition.

## References

1. Smits THM, Arend LNVS, Cardew S, Tång-Hallbäck E, Mira MT, Moore ERB, et al. Resolving taxonomic confusion: establishing the genus Phytobacter on the list of clinically relevant Enterobacteriaceae. European Journal of Clinical Microbiology & Infectious Diseases. 2022 Apr 15;41(4):547–58.

2. Medina-Cordoba LK, Chande AT, Rishishwar L, Mayer LW, Valderrama-Aguirre LC, Valderrama-Aguirre A, et al. Genomic characterization and computational phenotyping of nitrogen-fixing bacteria isolated from Colombian sugarcane fields. Sci Rep. 2021 Apr 28;11(1):9187.

3. Madhaiyan M, Saravanan VS, Blom J, Smits THM, Rezzonico F, Kim SJ, et al. Phytobacter palmae sp. nov., a novel endophytic, N2 fixing, plant growth promoting Gammaproteobacterium isolated from oil palm (Elaeis guineensis Jacq.). Int J Syst Evol Microbiol. 2020 Feb 1;70(2):841–8.

4. Zhang GX, Peng GX, Wang ET, Yan H, Yuan QH, Zhang W, et al. Diverse endophytic nitrogen-fixing bacteria isolated from wild rice Oryza rufipogon and description of Phytobacter diazotrophicus gen. nov. sp. nov. Arch Microbiol. 2008 May 4;189(5):431–9.

5. Salha Y, Sudalaimuthuasari N, Kundu B, AlMaskari RS, Alkaabi AS, Hazzouri KM, et al. Complete Genome Sequence of Phytobacter diazotrophicus Strain UAEU22, a Plant Growth-Promoting Bacterium Isolated from the Date Palm Rhizosphere. Microbiol Resour Announc. 2020 Jun 18;9(25).

6. Thele R, Gumpert H, Christensen LB, Worning P, Schønning K, Westh H, et al. Draft genome sequence of a Kluyvera intermedia isolate from a patient with a pancreatic abscess. J Glob Antimicrob Resist. 2017 Sep;10:1–2.

7. Choice S, Sherman A, Holder K, Harrington E. Gram-negative sepsis caused by a rare pathogen Phytobacter ursingii. BMJ Case Rep. 2024 Apr 16;17(4):e258384.

8. Lin J, Wu J, Gong L, Li X, Wang G. Sepsis caused by Phytobacter diazotrophicus complicated with galactosemia type 1 in China: a case report. BMC Infect Dis. 2024 Jun 19;24(1):599.

9. Pillonetto M, Arend L, Gomes SMT, Oliveira MAA, Timm LN, Martins AF, et al. Molecular investigation of isolates from a multistate polymicrobial outbreak associated with contaminated total parenteral nutrition in Brazil. BMC Infect Dis. 2018 Dec 13;18(1):397.

10. Rezzonico F, Smits THM, Duffy B. Misidentification slanders Pantoea agglomerans as a serial killer. Journal of Hospital Infection. 2012 Jun;81(2):137–9.

11. Hon P, Ko KKK, Zhong JCW, D. PP, Smits THM, Low J, et al. Genomic Identification of Two Phytobacter diazotrophicus Isolates from a Neonatal Intensive Care Unit in Singapore. Microbiol Resour Announc. 2023 Jun 20;12(6).

12. Pillonetto M, Arend LN, Faoro H, D’Espindula HRS, Blom J, Smits THM, et al. Emended description of the genus Phytobacter, its type species Phytobacter diazotrophicus (Zhang 2008) and description of Phytobacter ursingii sp. nov. Int J Syst Evol Microbiol. 2018 Jan 1;68(1):176–84.

13. Dal Lin A, Kulek DO, Gonçalves GA, Kraft L, Neto JFC, Vizentainer G, et al. Building of a new Spectra for the identification of Phytobacter spp., an emerging Enterobacterales, using MALDI Biotyper. Microbiol Spectr. 2024 Oct 3;12(10).

14. Michel IR, Kulek D, Arend LNVS, Pillonetto M, Smits THM, Rezzonico F. Development of two quantitative PCR assays for the detection of emerging opportunistic human pathogens belonging to the genus Phytobacter in routine diagnostics. Diagn Microbiol Infect Dis. 2024 Dec;110(4):116556.

15. Yaikhan T, Suwannasin S, Singkhamanan K, Chusri S, Pomwised R, Wonglapsuwan M, et al. Genomic Characterization of Multidrug-Resistant Enterobacteriaceae Clinical Isolates from Southern Thailand Hospitals: Unraveling Antimicrobial Resistance and Virulence Mechanisms. Antibiotics. 2024 Jun 6;13(6):531.

16. Huang Z, Zhang G, Zheng Z, Lou X, Cao F, Zeng L, et al. Genomic insights into the evolution, pathogenicity, and extensively drug-resistance of emerging pathogens Kluyvera and Phytobacter. Front Cell Infect Microbiol. 2024 Mar 21;14.

17. Almuzara M, Cittadini R, Traglia G, Haim MS, De Belder D, Alvarez C, et al. Phytobacter spp: the emergence of a new genus of healthcare-associated Enterobacterales encoding carbapenemases in Argentina: a case series. Infection Prevention in Practice. 2024 Sep;6(3):100379.

18. Kubota H, Nakayama T, Ariyoshi T, Uehara S, Uchitani Y, Tsuchida S, et al. Emergence of Phytobacter diazotrophicus carrying an IncA/C 2 plasmid harboring bla NDM-1 in Tokyo, Japan. mSphere. 2023 Aug 24;8(4).

19. Ma Y, Yao R, Li Y, Wu X, Li S, An Q. Proposal for Unification of the Genus Metakosakonia and the Genus Phytobacter to a Single Genus Phytobacter and Reclassification of Metakosakonia massiliensis as Phytobacter massiliensis comb. nov. Curr Microbiol. 2020 Aug 30;77(8):1945–54.

20. Adler A, Ayala-Montaño S, Assous M V., Geffen Y, Reuter S. Dissemination of a IncHI2A plasmid co-harboring the mcr-9 and blaNDM-1 genes in Israeli hospitals. Ann Clin Microbiol Antimicrob. 2025 Aug 20;24(1):45.

21. Fu Y, Morris FC, Pereira SC, Kostoulias X, Jiang Y, Vidor C, et al. Mechanisms of blaIMP-4 dissemination across diverse carbapenem-resistant clinical isolates. J Glob Antimicrob Resist. 2025 Mar;41:189–94.

22. Adler A, Ayala-Montaño S, Assous M V., Geffen Y, Reuter S. Dissemination of a IncHI2A plasmid co-harboring the mcr-9 and blaNDM-1 genes in Israeli hospitals. Ann Clin Microbiol Antimicrob. 2025 Aug 20;24(1):45.

23. Kovarova A, Amadasun M, Hooban B, McDonagh F, Tumeo A, Ryan K, et al. Characterisation of Citrobacter freundii and Enterobacter cloacae complex isolates co-carrying blaNDM-1 and mcr-9 from three hospitals. J Glob Antimicrob Resist. 2025 Sep;44:226–33.

24. Wright H, Bonomo RA, Paterson DL. New agents for the treatment of infections with Gram-negative bacteria: restoring the miracle or false dawn? Clinical Microbiology and Infection. 2017 Oct;23(10):704–12.

25. Cahill N, Hooban B, Fitzhenry K, Joyce A, O’Connor L, Miliotis G, et al. First reported detection of the mobile colistin resistance genes, mcr-8 and mcr-9, in the Irish environment. Science of The Total Environment. 2023 Jun;876:162649.

26. Tyson GH, Li C, Hsu CH, Ayers S, Borenstein S, Mukherjee S, et al. The mcr-9 Gene of Salmonella and Escherichia coli Is Not Associated with Colistin Resistance in the United States. Antimicrob Agents Chemother. 2020 Jul 22;64(8).

27. Dolejska M, Masarikova M, Dobiasova H, Jamborova I, Karpiskova R, Havlicek M, et al. High prevalence of Salmonella and IMP-4-producing Enterobacteriaceae in the silver gull on Five Islands, Australia. Journal of Antimicrobial Chemotherapy. 2016 Jan;71(1):63–70.

28. Peleg AY, Franklin C, Bell JM, Spelman DW. Dissemination of the Metallo--Lactamase Gene blaIMP-4 among Gram-Negative Pathogens in a Clinical Setting in Australia. Clinical Infectious Diseases. 2005 Dec 1;41(11):1549–56.

29. Nakano A, Nakano R, Suzuki Y, Saito K, Kasahara K, Endo S, et al. Rapid Identification of bla IMP-1 and bla IMP-6 by Multiplex Amplification Refractory Mutation System PCR. Ann Lab Med. 2018 Jul 28;38(4):378–80.

30. Harino T, Kayama S, Kuwahara R, Kashiyama S, Shigemoto N, Onodera M, et al. Meropenem Resistance in Imipenem-Susceptible Meropenem-Resistant Klebsiella pneumoniae Isolates Not Detected by Rapid Automated Testing Systems. J Clin Microbiol. 2013 Aug;51(8):2735–8.

31. Villa L, García-Fernández A, Fortini D, Carattoli A. Replicon sequence typing of IncF plasmids carrying virulence and resistance determinants. Journal of Antimicrobial Chemotherapy. 2010 Dec;65(12):2518–29.

32. Yang X, Heng H, Zhang H, Peng M, Chan EWC, Shum HP, et al. IncFIBK/FIIK conjugative iuc3-carrying virulence plasmids of clinical hypervirulent Klebsiella pneumoniae are multi-drug resistant. Microbiol Res. 2025 Nov;300:128288.

33. Wang X, Zhao J, Ji F, Chang H, Qin J, Zhang C, et al. Multiple-Replicon Resistance Plasmids of Klebsiella Mediate Extensive Dissemination of Antimicrobial Genes. Front Microbiol. 2021 Oct 27;12.

34. Chen S. Ultrafast one-pass FASTQ data preprocessing, quality control, and deduplication using fastp. iMeta. 2023 May 8;2(2).

35. Parks DH, Imelfort M, Skennerton CT, Hugenholtz P, Tyson GW. CheckM: assessing the quality of microbial genomes recovered from isolates, single cells, and metagenomes. Genome Res. 2015 Jul;25(7):1043–55.

36. Meier-Kolthoff JP, Carbasse JS, Peinado-Olarte RL, Göker M. TYGS and LPSN: a database tandem for fast and reliable genome-based classification and nomenclature of prokaryotes. Nucleic Acids Res. 2022 Jan 7;50(D1):D801–7.

37. Seemann T. Prokka: rapid prokaryotic genome annotation. Bioinformatics. 2014 Jul 15;30(14):2068–9.

38. Page AJ, Cummins CA, Hunt M, Wong VK, Reuter S, Holden MTG, et al. Roary: rapid large-scale prokaryote pan genome analysis. Bioinformatics. 2015 Nov 15;31(22):3691–3.

39. Stamatakis A. RAxML version 8: a tool for phylogenetic analysis and post-analysis of large phylogenies. Bioinformatics. 2014 May 1;30(9):1312–3.

40. Huerta-Cepas J, Serra F, Bork P. ETE 3: Reconstruction, Analysis, and Visualization of Phylogenomic Data. Mol Biol Evol. 2016 Jun;33(6):1635–8.

41. Robertson J, Nash JHE. MOB-suite: software tools for clustering, reconstruction and typing of plasmids from draft assemblies. Microb Genom. 2018 Aug 1;4(8).

42. Alcock BP, Huynh W, Chalil R, Smith KW, Raphenya AR, Wlodarski MA, et al. CARD 2023: expanded curation, support for machine learning, and resistome prediction at the Comprehensive Antibiotic Resistance Database. Nucleic Acids Res. 2023 Jan 6;51(D1):D690–9.

